# DeepSAP: Improved RNA-Seq Alignment by Integrating Transcriptome Guidance with Transformer-Based Splice Junction Scoring

**DOI:** 10.1101/2025.04.23.650072

**Authors:** Fadel Berakdar, Thomas D. Wu, Tong Zhu, Mehrzad Samadi, Pankaj Vats

## Abstract

Advancements in high-throughput sequencing have revolutionized the field of transcriptomics, providing unprecedented insights into gene expression, splicing events, and fusions. Despite these advancements, the analysis of RNA-seq data remains challenging due to the presence of complex splice junctions, multi-mapped reads, and chimeric events. In this study, we present DeepSAP, an innovative approach that improves the accuracy of RNA-seq alignment by integrating Transcriptome-Guided Genomic Alignment, as implemented in GSNAP, with improved splice junction scoring, predicted by a transformer-based deep learning model. Our work demonstrates synergy between these methods, resulting in enhanced detection of splice junctions, identification of indels, and resolution of complex splicing patterns. On a standard benchmark of human simulated datasets, DeepSAP achieves the highest mean F1 score (0.971) for splice junction detection, outperforming DRAGEN (0.933), novoSplice (0.914), STAR (0.821), HISAT2 (0.662), and Subjunc (0.770). By integrating the unique capabilities of transcriptome-guided alignment and large language models, our splice junction scoring approach captures intricate sequence patterns surrounding splice donor and acceptor sites, providing significant advancement in RNA-seq data analysis.

## Introduction

High-throughput sequencing has revolutionized our understanding of molecular biology, with RNA-seq enabling a comprehensive exploration of the transcriptome. RNA-seq enables researchers to quantify differential gene expression under various conditions or treatments, as well as identify both annotated and novel transcripts. These transcripts can reveal gene fusions, alternative splicing events, and non-coding RNA that may have been overlooked by previous methodologies. However, achieving such insights into gene regulation and function depends critically on accurate read alignment. RNA-seq protocols produce short fragments with read lengths of 100–200 base pairs, either from both ends (paired-end) or from one end (single-end) of a cDNA fragment. Aligning short reads to the genome is challenging due to sequencing errors, as well as issues unique to RNA-seq including intronic regions, alternative splicing, incomplete reference transcriptomes, non-coding RNA, pseudogenes, ambiguous splicing patterns, and RNA editing. In particular, as applications of RNA-seq move toward the analysis of individual transcriptomes, which have shown surprising variability(Glinos et al. [2022]), it is becoming increasingly important to handle such variations accurately.

The inherent complexities in RNA-seq data necessitate the use of specialized alignment algorithms and tools designed to address their unique characteristics, making RNA-seq alignment significantly more challenging than DNA-seq. Although current RNA-seq alignment programs handle most reads effectively, biologically significant subsets remain problematic, particularly reads containing insertions/deletions, reads spanning cryptic splice junctions, and those originating from genomic regions with SNPs or mutations. Another major issue is multi-mapped reads, as these sequences align to multiple genomic locations with similar confidence scores. To ensure accurate results, aligners must consider all possible genomic locations during the alignment process.

To address these challenges, a variety of splice-aware alignment tools have been developed over the years. The first category includes alignment-based RNA-seq aligners like GSNAP(Wu et al. [2016]), STAR(Dobin et al. [2013]) and HISAT2(Kim et al. [2019]). These aligners perform a complete alignment between the read and the genomic reference, producing a BAM file as output. They offer high precision in base-level alignment and junction detection, making them ideal for comprehensive RNA-seq analysis. In contrast, tools such as Salmon(Patro et al. [2017]) and Kallisto(Bray et al. [2016]) employ alignment-free or pseudo-alignment methods. For example, quasi-mapping in Salmon and pseudo-alignment in Kallisto function by matching k-mers of reads to a transcriptomic reference without performing full alignments. These approaches significantly accelerate the process and can generally provide high accuracy in transcript quantification, making them suitable for estimating transcript abundance. However, these methods may be less effective in detecting mismatches, insertions and deletions (indels), novel isoforms, or alternative splicing events, and may underperform compared with traditional alignment tools in small low-abundance genes or for novel splicing events and non-coding RNA.

The performance of RNA-seq alignment tools has been evaluated in previously published benchmarking studies. Notably, a study conducted by Baruzzo et al.(Baruzzo et al. [2017]) performed an exhaustive benchmarking analysis of 14 widely used RNA-seq aligners utilizing simulated datasets derived from annotated transcriptomes of two different species, human and malaria. The study evaluated the performance of each tool with multiple criteria, including base-level accuracy, read-level alignment, and splice junction detection. The datasets in the study were designed to simulate different levels of complexity, emulating real biological variations, sequencing errors, and were categorized as T1, T2, and T3 datasets for both human and malaria. The results revealed significant disparities in the performance of aligners, particularly in their proficiency at detecting insertions and deletions, mismatches, and splice junctions. In the study conducted by Baruzzo et al., as well as in a separate study by Grant et al.(Grant et al. [2011]), GSNAP exhibited superior performance at both the read and base levels when compared to other aligners. However, despite its strong performance, GSNAP, along with other aligners, demonstrated areas for improvement, particularly at the T3 level of alignment complexity.

To further enhance alignment performance, we have developed DeepSAP (Deep Splice Alignment Program), a novel approach that integrates a new feature called Transcriptome-Guided Genomic Alignment (TGGA) in GSNAP, complemented by advanced deep learning techniques, such as fine-tuned transformer models (Figure 1). The TGGA feature leverages a given transcriptome (such as those used by alignment-free methods) to allow for a relatively straightforward alignment of reads to known transcripts, while retaining the capacity for full genomic alignment to accommodate novel splice phenomena. The concept of combining transcriptome and genome information for alignment was explored previously in the RNA-seq Unified Mapper (RUM)(Grant et al. [2011]). However, successful implementation of the idea is challenging, which our work has attempted to address. Specifically, while aligning to a transcriptome is much simpler than aligning to a genome because intron gaps do not need to be considered, a read still potentially contains mismatches and indels compared with its transcript of origin. Therefore, we have found that several methods are required to perform transcriptome alignment accurately and efficiently. In our implementation, TGGA GSNAP employs three methods in successive order: exact search, prevalence search, and extension search, until a satisfactory alignment is obtained (Figure 1). The criterion for a satisfactory alignment is another issue in implementation. For paired-end reads, TGGA GSNAP searches for a concordant alignment of the two ends to the same transcript; for single-end reads, TGGA GSNAP looks for an alignment with 95% or more of the nucleotides in the read matching the transcript. If a satisfactory transcriptome alignment cannot be found, then TGGA GSNAP attempts to find an alignment to the genome using its standard methods, which are already among the most accurate, thereby enhancing the overall accuracy of TGGA.

**Figure 1:**
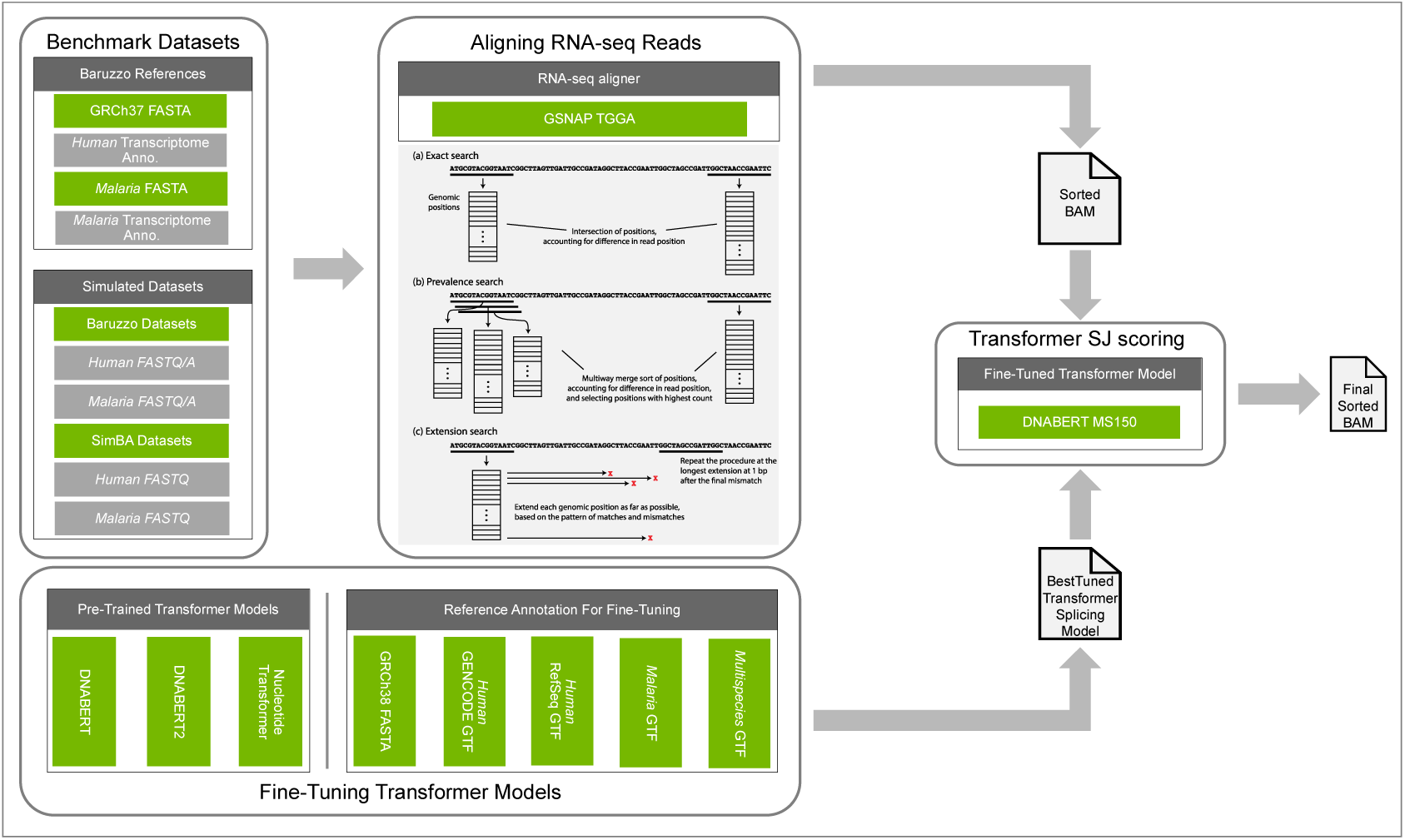
Overview of the DeepSAP workflow. DeepSAP utilizes the TGGA GSNAP aligner initially to align short-read RNA-seq data. The TGGA aligner tries three methods for transcriptome alignment, exact search, prevalence search, and extension search, to find a satisfactory solution, and otherwise resorts to genomic alignment. In the next phase, it incorporates the fine-tuned DNABERT MS150 transformer model to enhance splice junction detection. This second stage analyzes all alignments generated by TGGA GSNAP for each query read, recalibrates the MAPQ (mapping quality) scores for multi-mapped reads and applies soft clipping for splice junctions that exhibit either low transformer scores or flanking bases with suboptimal Phred quality scores.

Building upon the foundation of alignment using TGGA, we have also integrated state-of-the-art LLMs, built on transformer-based architectures, to improve RNA-seq alignment accuracy through enhanced scoring of splice junctions. This novel methodology leverages transformers(Vaswani et al. [2017]), which excel at capturing long-range dependencies and complex sequence patterns. Transformers are specifically designed to process and understand sequences by focusing on relationships between elements using a mechanism called “attention.” This enables them to effectively capture context and meaning within data. LLMs, such as GPT and BERT, are large-scale implementations of transformers trained on massive datasets. While originally developed for natural language processing tasks like generating human-like text, answering questions, and translating languages, transformers have also shown tremendous potential in genomics. Models such as DNABERT(Ji et al. [2021]), DNABERT2(Zhou et al. [2024]), and Nucleotide Transformer(Dalla-Torre et al. [2023]) highlight the adaptability of transformers to biological sequence analysis.

Other deep learning methods have been applied to improve the accuracy of splice site prediction, starting with SpliceAI(Jaganathan et al. [2019]), which used a deep convolutional neural networks architecture, whereas another deep learning method Splam(Chao et al. [2024]) uses deep residual neural networks. Many researchers have shown that transformer-based models can outperform neural network models, notable examples include SpliceTransformer(You et al. [2024]), Transformer-45k(Jónsson et al. [2024]), and TrASPr(Wu et al. [2025]). These models have generally been used to evaluate splice junctions inferred from an entire set of RNA-seq alignments, or to choose among different alternative splice isoforms, especially in a tissue-specific context. In contrast, our work focuses on improving individual read alignments, which serve as the foundation for all subsequent analyses. Improving splice site scoring at the alignment stage is expected to be more effective than at later analytical stages, as downstream analyses often rely on the number of reads supporting a specific splice site or indel. Furthermore, the extent to which state-of-the-art machine learning techniques can impact RNA-seq analysis remains uncertain. In this study, we perform extensive benchmarking and comparisons with existing aligners to quantify the utility of such techniques for RNA-seq alignment.

Another goal of our study is to develop a practical tool that leverages transformer models to enhance splice junction detection in RNA-seq alignments by applying transformer-based splice junction scoring in a post-alignment step. To optimize these models for RNA-seq data, we fine-tuned pre-trained transformers that were initially trained on extensive genomic datasets. This fine-tuning process has enabled the models to accurately predict splice sites and improve the results of RNA-seq alignment. Although our transformer-based workflow could process outputs from any aligner that provides the XS SAM tag, it performs best when paired with highly sensitive aligners. The need for sensitivity motivated us to use GSNAP, which excels at detecting complex splice junctions and indels, especially those near splice sites. In this study, we observed that the combination of TGGA GSNAP and Transformer guided splice junction detection achieves high accuracy in RNA-seq alignment, as demonstrated by our benchmarking results.

## Results

### Fine-Tuning of Transformer Models for Splice Site Scoring

To optimize transformer models for RNA-seq data, we fine-tuned pre-trained transformers that were initially trained on extensive genomic datasets. We evaluated various transformer models, including DNABERT 6-mer(Ji et al. [2021]), DNABERT2(Zhou et al. [2024]), and the 2.5B multi-species Nucleotide Transformer(Dalla-Torre et al. [2023]).

These models were fine-tuned on labeled donor and acceptor site sequences, as well as negative examples. Negative examples were randomly generated from the genome by selecting two loci based on the following criteria: they do not overlap with donor or acceptor sites, are located within a predefined distance range from each other, maintain a predefined ratio of splicing signals similar to those in the true examples, and match the number of negative examples with true examples. We used separate labels for donor, acceptor and negative examples. The motif sequences of the acceptor and donor sites were kept in the center as input for the transformer models. The input sequence lengths were fixed per dataset at 90, 150, 200, or 400 bases and generated using multiple sources: (i) GRCh38 NCBI RefSeq assembly (GCF_000001405.40), (ii) GRCh38 GENCODE release 44, (iii) malaria ASM276v2, and (iv) a multi-species dataset that included splice junctions from *Homo sapiens*, *Arabidopsis thaliana*, *Zea mays*, *Xenopus tropicalis*, *Saccharomyces cerevisiae*, *Drosophila melanogaster*, *Caenorhabditis elegans*, *Rattus norvegicus*, *Mus musculus*, *Plasmodium falciparum*, *Plasmodium vivax*, *Plasmodium berghei*, *Plasmodium knowlesi*.

The fine-tuning datasets could therefore be categorized based on species and sequence length: RefSeq90, RefSeq150, RefSeq200, and RefSeq400 for human RefSeq data; GENCODE150 and GENCODE400 for GENCODE data; Malaria90, Malaria150, Malaria200, and Malaria400 for malaria data; and MS150 for a multispecies dataset. Each dataset was randomly shuffled and split into training (70%), testing (15%), and validation (15%) sets, ensuring no data leakage between them. The validation set remained completely unseen during training and testing. The number of fine-tuning examples increased as sequence length decreased, given that the exon and intron lengths had to exceed half the window size. For instance, RefSeq90 included 630,424 sequences, whereas RefSeq400 contained 58,760. This extensive dataset facilitated the effective fine-tuning of transformer models for splice junction prediction.

We evaluated our fine-tuned models using the validation dataset to measure their effectiveness in identifying splice sites. Their performance was assessed using several metrics (Figure 2a): Loss function, which measures the deviation of the model’s predictions from the correct answers (lower loss is better); MCC (Matthews Correlation Coefficient), which indicates the overall accuracy of the model, even with imbalanced data; F1 Score, which balances Precision and Recall to show how well the model avoids false predictions; Recall, which indicates how many true splice sites the model detected; Accuracy, which represents the overall percentage of correct predictions; and Precision, which measures the proportion of correctly identified splice sites. These metrics helped us evaluate the model’s reliability in identifying complex splice patterns in genetic data.

**Figure 2:**
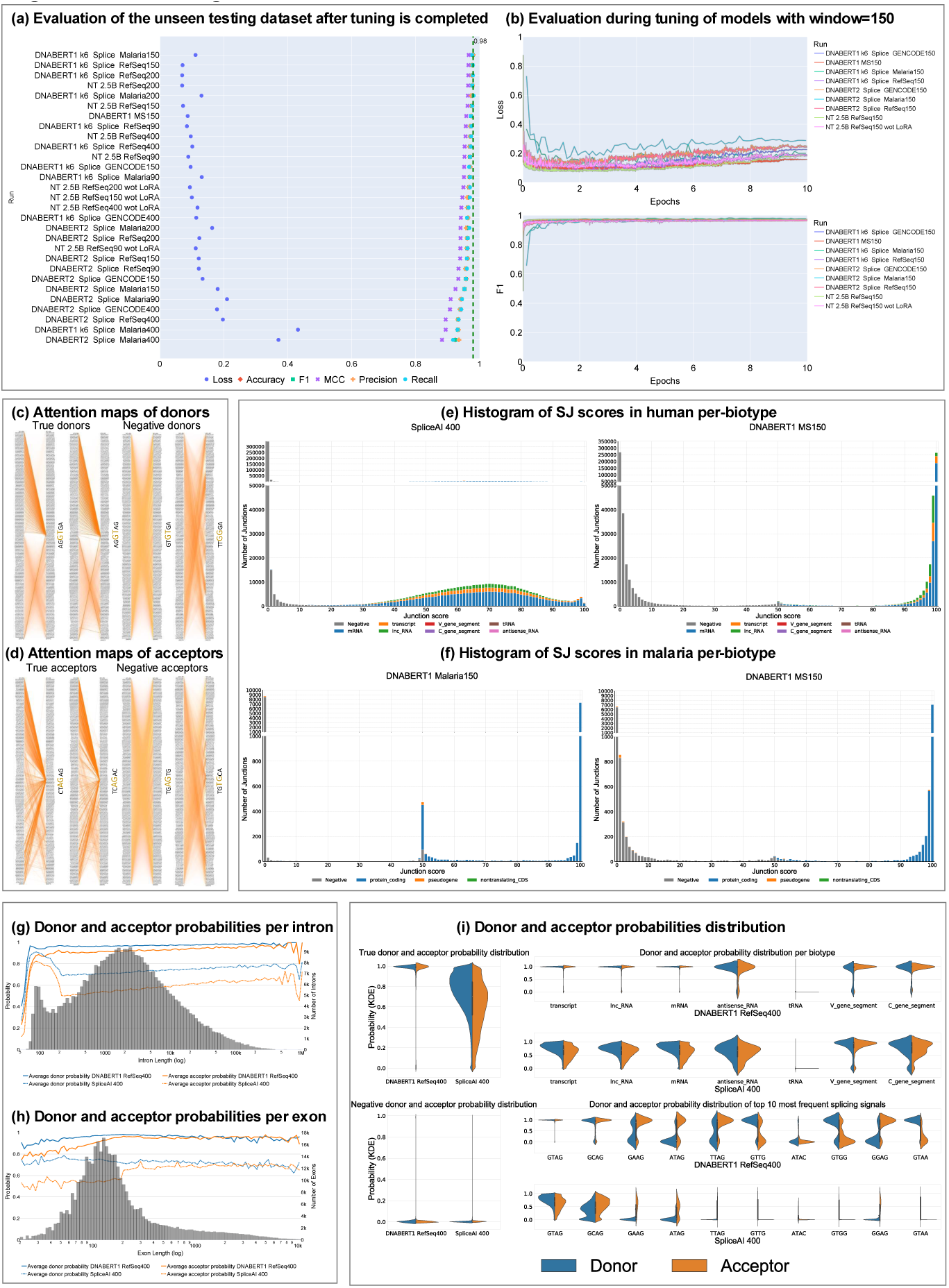
(a) Performance of fine-tuned models on the test dataset, ranked by MCC (Matthews Correlation Coefficient). (b) Loss and F1 score measured every 200 steps during fine-tuning. (c) Attention maps for true and negative donor splice sites generated by DNABERT MS150 representing head 12 of layer 12. (d) Attention maps for true and negative acceptor splice sites generated by DNABERT MS150 representing head 12 of layer 12. (e) Comparison of splice junction scores (average of donor and acceptor probabilities) between SpliceAI400 and DNABERT MS150 in human RefSeq splice junctions. (f) Comparison of splice junction scores between DNABERT Malaria150 and DNABERT MS150 in malaria splice junctions. (g) Donor and acceptor scores across intron lengths for DNABERT RefSeq400 and SpliceAI400. (h) Donor and acceptor scores across exon lengths for DNABERT RefSeq400 and SpliceAI400. (i) Distribution of donor and acceptor probabilities across various biotypes and splicing signals in human RefSeq splice junctions for DNABERT RefSeq400 and SpliceAI400.

Our results demonstrated improved performance when using RefSeq datasets, achieving the highest Matthews Correlation Coefficient (MCC) across all fine-tuned transformer models (Figure 2a). Notably, the DNABERT model consistently outperformed DNABERT2, regardless of sequence length or genome annotation. For instance, the MCC measurements showed: DNABERT RefSeq150 at 0.965, DNABERT MS150 at 0.958, DNABERT2 RefSeq150 at 0.934, and NT at 0.964. Furthermore, the species-specific DNABERT fine-tuned models achieved higher MCC scores than DNABERT MS150 for both malaria and human datasets. Our analysis revealed that the multi-species DNABERT MS150 demonstrates a stronger ability to distinguish correct splice junctions, clustering them closer to a score of 100 versus negative splice junction examples close to 0 (Figure 2e). This advantage was particularly evident in the malaria datasets, where the multi-species model outperformed the malaria-specific model in differentiating between true donor and acceptor sites (Figure 2f).

Moreover, attention maps generated by the DNABERT MS150 model reveal that at true donor and acceptor splice sites, which contain canonical GT or AG dinucleotides, the model assigns high attention weights to 6-mers up to 75 base pairs upstream of donor splice sites and both upstream and downstream of acceptor splice sites. These maps suggest that, among potential splice sites with canonical dinucleotides, exon content is highly critical for predicting donor splices, perhaps due to differing codon usage in coding regions and introns. For acceptor splice sites, the upstream region may correspond to sequences critical for the spliceosome’s function, such as the branch point sequence. In contrast, the attention weights around negative examples are significantly lower than those of positive examples (Figure 2c-d), implying that the local context around the splice site plays an important part in prediction. These attention maps highlight the advantage of long-range models that transformers can analyze, compared with local models such as MaxEnt(Yeo and Burge [2003]), which look only at the 3 to 9 base pairs surrounding the splice site. This finding is consistent with other studies(Jaganathan et al. [2019]) that have found that using relatively long regions is favorable for splice site prediction.

While exon and intron lengths are not directly used as inputs to the transformer scoring function, we observed a notable relationship between predicted splice site probabilities and these lengths (Figure 2g-h). Specifically, intron lengths exhibit a bimodal distribution, whereas exon lengths follow a unimodal distribution, aligning with established biological patterns. The bimodal distribution of intron lengths is well-documented in the literature and has been attributed to evolutionary mechanisms that favor distinct classes of introns(Zhang et al. [2016]). Furthermore, DNABERT’s performance on the RefSeq400 dataset demonstrates its ability to handle complex splicing motifs more effectively than SpliceAI400 (Figure 2i). Notably, DNABERT achieves superior accuracy in predicting rarer biotypes and splicing signals across different species. These findings underscore DNABERT’s adaptability and robustness in capturing intricate splicing patterns.

### Benchmarking RNA-seq Aligners

We evaluated DeepSAP’s performance using the Baruzzo(Baruzzo et al. [2017]) and SimBA(Audoux et al. [2017]) simulated datasets against several state-of-the-art RNA-seq aligners, including Illumina’s DRAGEN™ v4.0.3, novoSplice v0.8.4(Berakdar et al. [2019]), STAR v2.7.10a(Dobin et al. [2013]), HISAT2 v2.2.1(Kim et al. [2019]) and Subjunc v2.0.1(Liao et al. [2013]). All aligners were benchmarked using their default settings.

In our benchmark, the DeepSAP workflow, powered by TGGA GSNAP and the DNABERT MS150 transformer model, demonstrated outstanding performance across all evaluated metrics, consistently achieving best results (Figure 3a). Key metrics included recall and precision at read and base levels. Read-level metrics were satisfied if at least one base in a read was aligned correctly, whereas base-level metrics assessed the accuracy of calling each nucleotide position. DeepSAP ranked top in correctly aligning bases across Baruzzo et al. benchmark datasets (Figure 3b).

**Figure 3:**
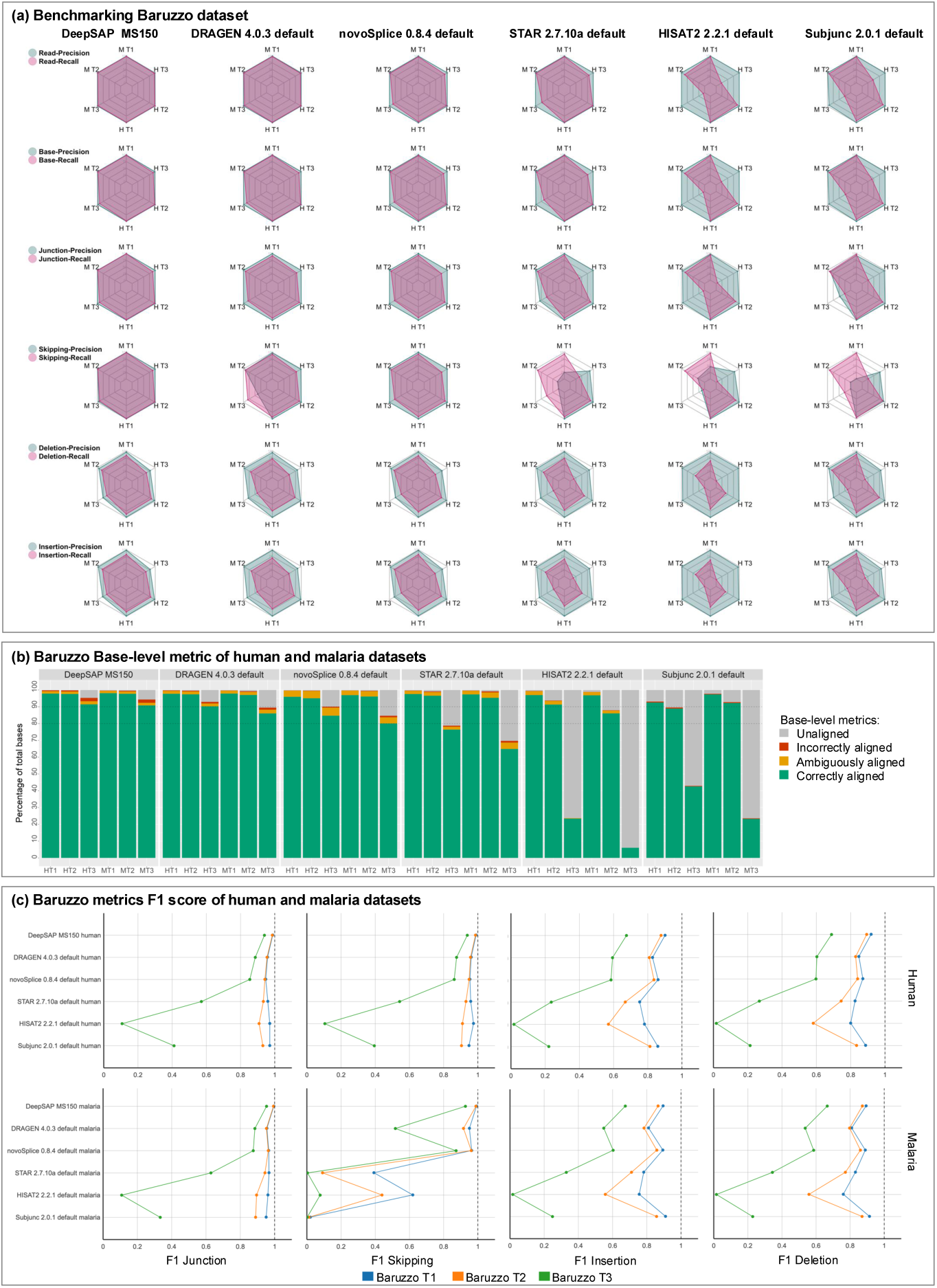
(a) Overview of the benchmarking of the Baruzzo human (HT1, HT2, HT3), and malaria (MT1, MT2, MT3) datasets. DeepSAP and DRAGEN excel at the read and base level, with DeepSAP performing exceptionally well in the skipping metric, which assesses the detection of junctions per spliced read, unlike the junction metric, which measures junctions as single events. (b) Base-level metric in human and malaria datasets, DeepSAP has the highest aligned bases. (c) Scatter plot of F1 scores for Baruzzo metrics in detecting junctions, skipping, insertion, and deletion events, highlighting DeepSAP’s consistent top performance, with DRAGEN and novoSplice performing closely.

In addition to demonstrating notable accuracy in read and base-level metrics, DeepSAP excelled in detecting splice junctions, as evidenced by the junction and skipping metrics, measured through recall and precision (Figure 3a), as well as their F1 scores (Figure 3c). The junction metrics evaluated splice junctions on an individual basis for each read, whereas the skipping metrics assessed splice junction accuracy across the entire dataset, determining whether a splice junction was correctly or incorrectly identified by any read within the dataset.

Moreover, DeepSAP achieved high scores in detecting indels, based on both recall and precision metrics (Figure 3a) and F1 scores (Figure 3c, Supplemental Table.1). Our workflow demonstrated a strong capability to handle reads with complex insertions and deletions, particularly from T3 datasets. Furthermore, in another independent benchmarking dataset generated utilizing SimBA(Audoux et al. [2017]), which exhibited extremely high levels of sequence variations compared with the reference genome, DeepSAP workflow consistently maintained high performance (Supplemental Fig. S3).

### Detection of Complex Splice Junctions

We evaluated the performance of various aligners in detecting splice junctions using both simulated and real datasets. Scatterplots comparing splice junction counts with the ground truth from the Baruzzo simulated datasets (Figure 4a) revealed significant variability among aligners, particularly in the Baruzzo T3 dataset. Among the evaluated methods, the DeepSAP workflow demonstrated the closest agreement to the ground truth, as evidenced by the Theil-Sen slope indicating strong concordance along the diagonal. This highlights DeepSAP’s superior ability to accurately capture linear trends between its splice junction counts and the ground truth. Additionally, scatterplots comparing aligners directly against each other (Figure 4b) showed that DeepSAP exhibited strong correlations with DRAGEN and novoSplice for both known and novel splice junction counts. Notably, DeepSAP identified a greater number of splice junctions compared to Subjunc, STAR, and HISAT2, as illustrated in (Figure 4a and Figure 4b). These results underscore DeepSAP’s robust performance in detecting splice junctions across diverse datasets.

**Figure 4:**
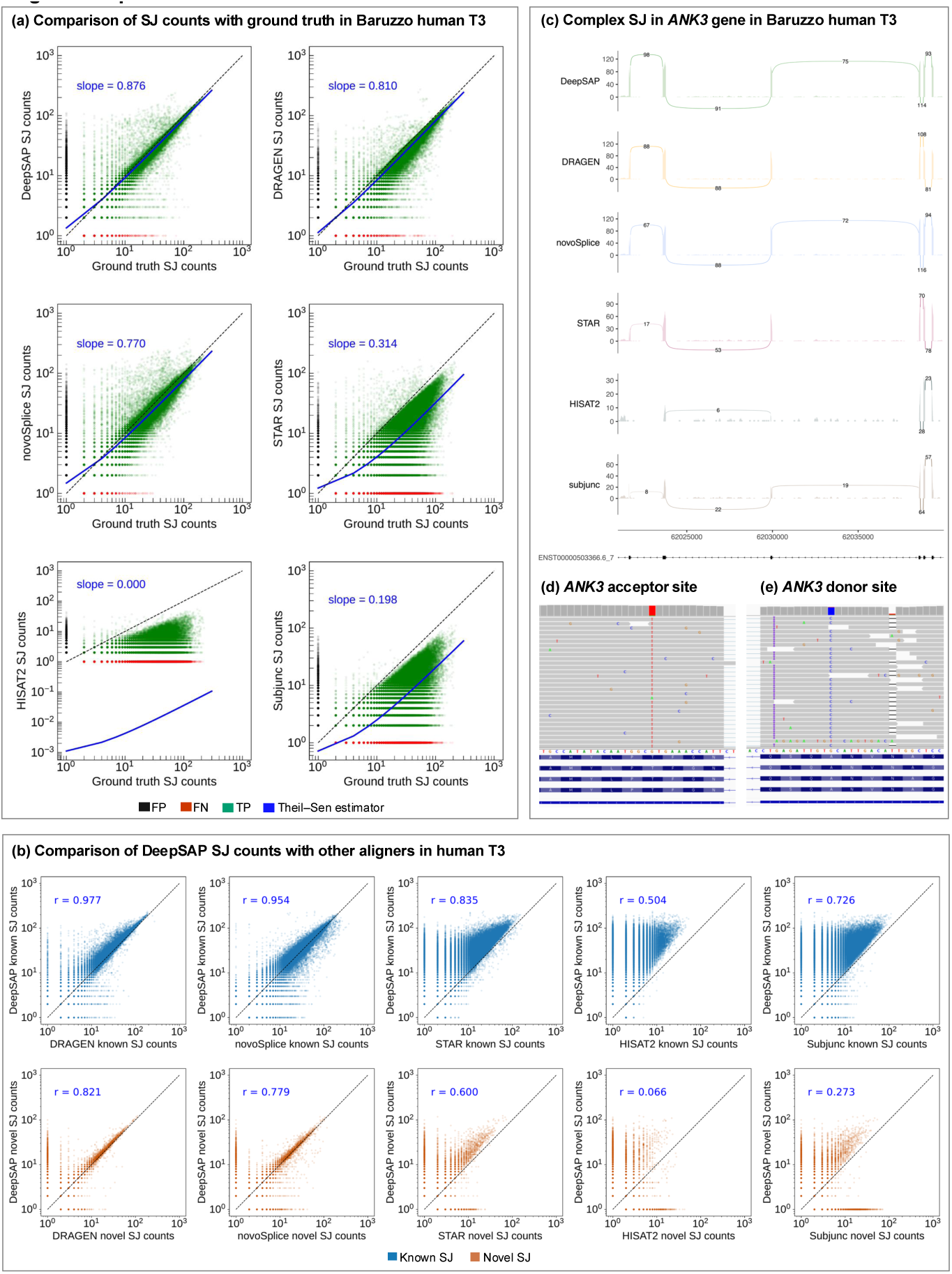
(a) Comparison of splice junction counts between the ground truth and each aligner in the Baruzzo human T3 dataset. The axes are plotted on a log scale, with 1 added to each count to avoid log(0) values. Theil-Sen slope was computed to estimate the linear trend between each aligner SJ counts and the ground truth. DeepSAP shows the best Theil-Sen slope in comparison with other aligners. (b) Pairwise comparison of splice junction counts between DeepSAP and the other aligners. The axes are plotted on a log scale, with 1 added to each count to avoid log(0) values with blue indicating known splice junctions, while orange representing novel junctions. DeepSAP and DRAGEN, as well as DeepSAP and novoSplice, show high Pearson correlation with significant p-values for known and novel junctions. (c) A splice junction in the *ANK3* gene from the Baruzzo human T3 dataset, detected by DeepSAP and novoSplice but missed by other aligners. (d) The acceptor site in the *ANK3* gene, showing a mismatch near the acceptor site. (e) An insertion and mismatch near the donor site of the *ANK3* gene.

DeepSAP demonstrates superior performance on T3 datasets, establishing its effectiveness for analyzing individual transcripts with complex splicing and indel variations. Our methodology achieves significantly higher T3 accuracy compared to previous benchmarks on these challenging datasets. For example, we highlight such an improvement in the *ANK3* gene (Figure 4c-e) which contains simulated mismatches and insertions near both donor and acceptor splice sites and was detected with the highest sensitivity by DeepSAP and novoSplice.

Similarly, scatterplots (Figure 5a) demonstrate a strong Pearson correlation with significant p-values when comparing DeepSAP to DRAGEN and novoSplice for both known and novel splice junctions in real datasets. For instance, in the publicly available pHGG sample SRR5280319, our workflow identified a novel splicing variant involving a retained intron in the *SMARCC2* gene (Figure 5b), which was missed by other aligners and not found in a previous survey of alternate splicing in high-grade gliomas using STAR(Siddaway et al. [2022]). This splice junction was characterized by multiple mismatches near its splice sites (Figure 5c-d), explaining why it was observed only by our workflow.

**Figure 5:**
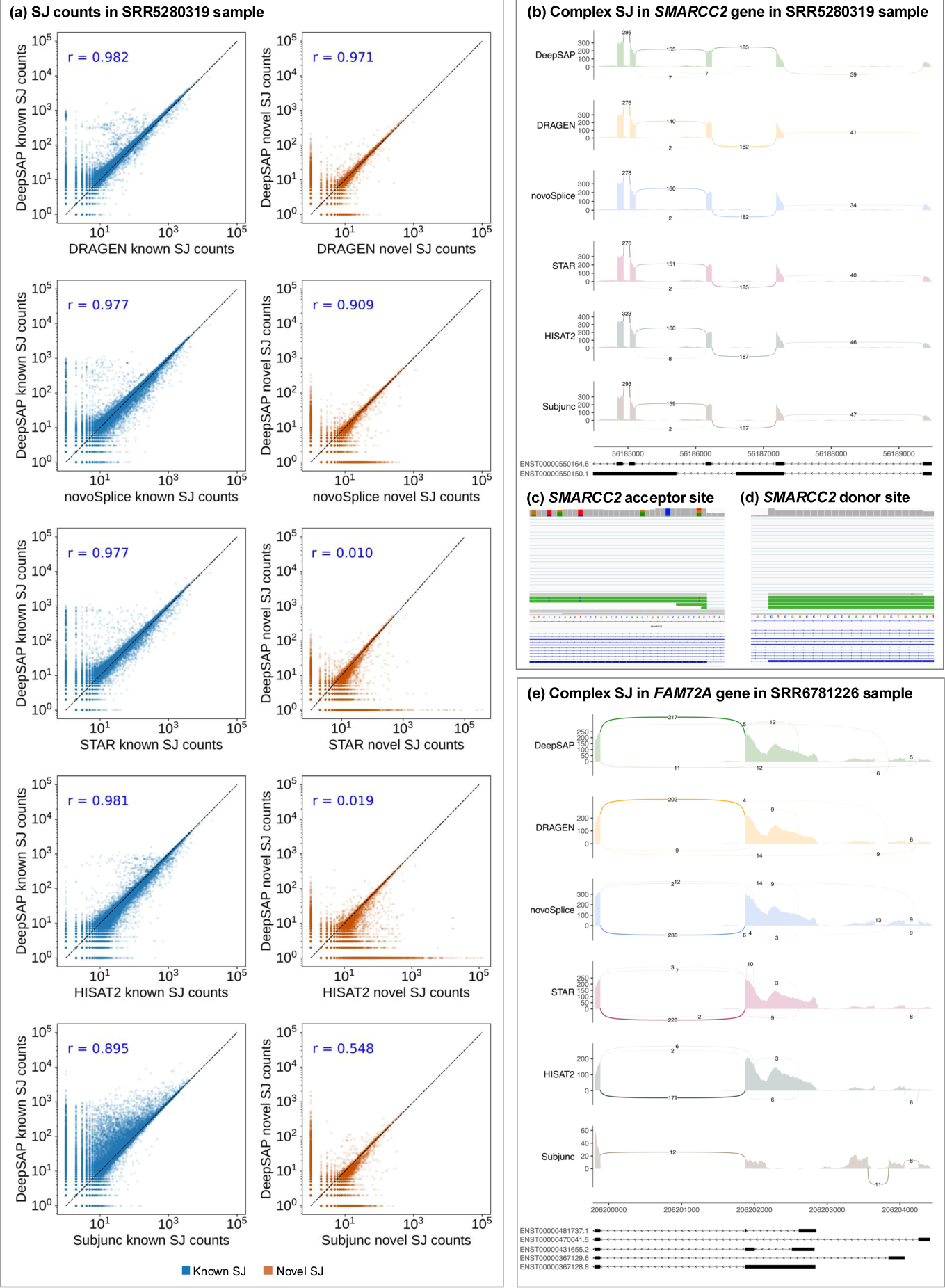
(a) Comparison of splice junctions between DeepSAP and other aligners in the SRR5280319 dataset from the pHGG study. The axes are plotted on a log scale, with 1 added to each count to avoid log(0) values with blue indicating known splice junctions, while orange representing novel junctions. DeepSAP and DRAGEN, as well as DeepSAP and novoSplice, show high Pearson correlation with significant p-values for known and novel junctions. (b) A splice junction in *SMARCC2* (SRR5280319), detected by DeepSAP but missed by other aligners, corresponds to an annotated transcript of the *SMARCC2* gene featuring a retained intron. (c) The acceptor site of the *SMARCC2* splice junction, showing multiple mismatches near the acceptor site. (d) The donor site of the *SMARCC2* splice junction, showing clean alignment. (e) Splice junction patterns in *FAM72A* gene (SRR6781226).

Another real-world example was observed in myelodysplastic syndrome sample SRR6781226 for the *FAM72A* gene (Figure 5e), which plays a crucial role in regulating DNA damage repair and maintaining genome stability(Feng et al. [2021]; Rogier et al. [2021]; Fu et al. [2023]). In our analysis, complex splice junctions supported by annotated ENCODE transcripts were identified by DeepSAP, DRAGEN, and novoSplice, but missed by other aligners. For instance, the splice junction in transcript ENST00000470041.5 was identified with coverage of 12, 14, and 14 reads by DeepSAP, DRAGEN, and novoSplice, respectively, while remaining undetected by other aligners. Similarly, the splice junction in transcript ENST00000367129.6 was detected with coverage of 5, 4, and 4 reads by DeepSAP, DRAGEN, and novoSplice, respectively, and was missed by other aligners. These findings emphasize the importance of alignment tools with enhanced sensitivity and precision. These tools are crucial for accurately detecting complex splice junction events, particularly in the context of understanding gene expression and splicing dynamics in cancer and other complex diseases.

### Detecting Insertions and Deletions Events

To evaluate the performance of different aligners in detecting indels, we conducted a comprehensive evaluation of indel detection using the Baruzzo et al. simulated datasets. Rates of true positive insertion results across aligners ranged from 64.34 to 77.42% in human T1 dataset, with DeepSAP, DRAGEN, and novoSplice having the largest percentages (Figure 6a). For the malaria T1 dataset, true positives ranged from 68.79 to 72.76%, with similar results across aligners. DeepSAP showed higher sensitivity overall, with fewer false negatives but more false positives. For human and malaria T2 datasets, rates of true positives were generally the same as for their corresponding T1, with the exception of STAR and HISAT2 in human T2, which showed decreases in true positives to 57.88% and 49.98%, respectively, and STAR in malaria T2, where rates fell to 55.94%. For the T3 dataset, identification of insertions was somewhat lower in all aligners compared with the T2 dataset, with dramatically worse results for STAR, which has a true positive rate of 20.86% in human T3. HISAT2 and Subjunc performed particularly poorly, with true positive rates near zero in both human and malaria T3 datasets. For deletion events, the performance of aligners in human T1, T2, and T3 datasets was comparable to their performance on the corresponding insertion events. In the malaria dataset, however, aligners demonstrated slightly lower accuracy for deletion events compared to insertion events, with true positive rates ranging from 59.66% to 65.98% across all aligners in T1 dataset and from 57.74% to 66.60% in T2 dataset. Notably, most aligners maintained robust performance in detecting deletion events in malaria T3 dataset, achieving true positive rates between 56.19% and 62.63% across all aligners, except HISAT2 and Subjunc, which exhibited significantly lower rates compared to their performance in T1 and T2 datasets.

**Figure 6:**
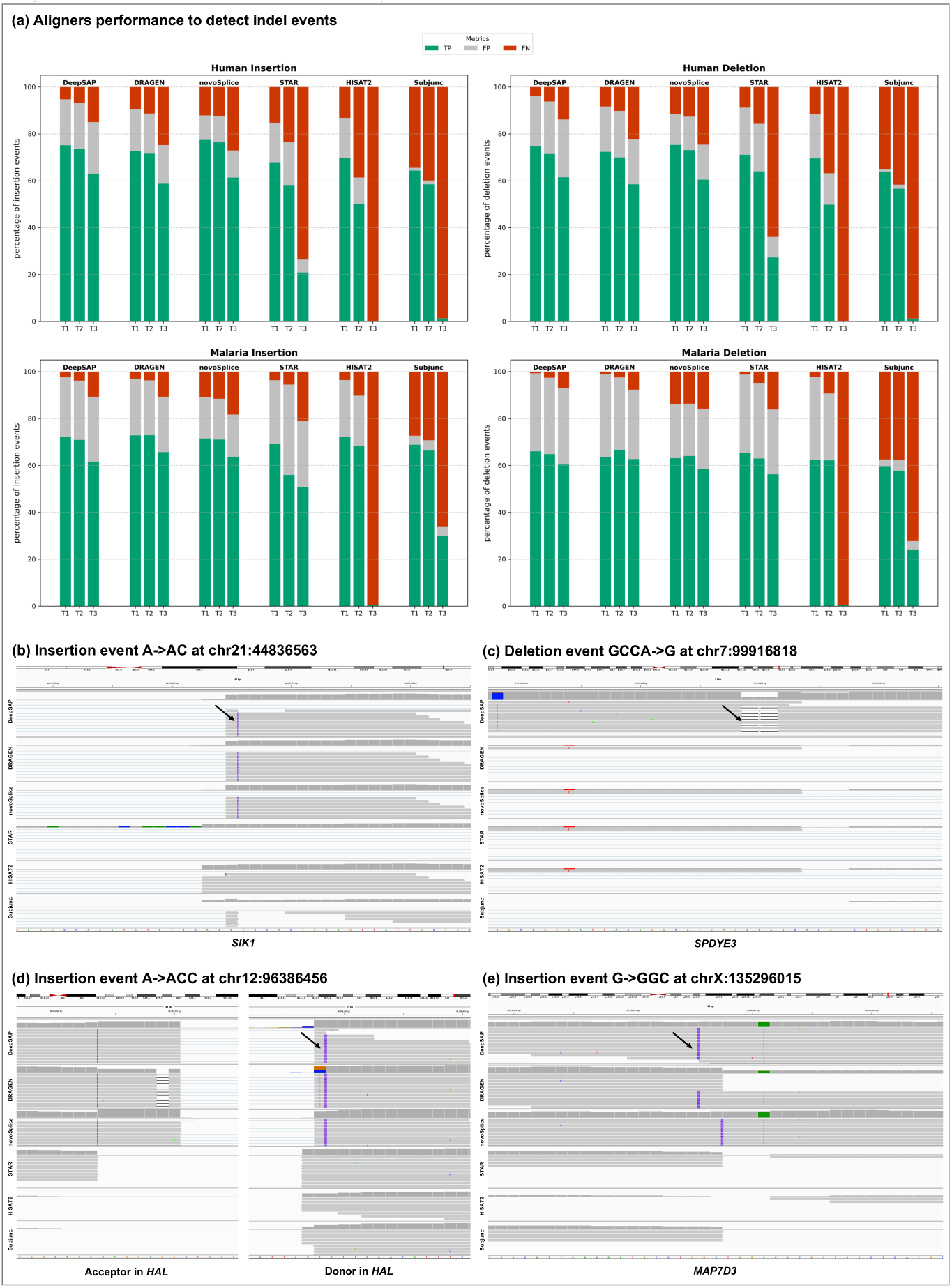
(a) The percentage of insertion and deletion events identified across the Baruzzo datasets. The bars are color-coded: green represents true positive events (correctly identified indels), red indicates false negatives (missed indels), and grey corresponds to false positives (incorrectly identified indels). (b) A true single-nucleotide insertion of C following an A at position chr21:44,836,563 in the *SIK1* gene detected by DeepSAP, novoSplice, and DRAGEN. (c) True deletion of three-base CCA following a G in the *SPDYE3* gene at position chr7:99,916,818 detected by DeepSAP only. (d) A true dinucleotide insertion of CC following an A in the *HAL* gene at position chr12:96,386,456 detected by DeepSAP and novoSplice. (e) A true dinucleotide insertion of GC after G, in the *MAP7D3* gene at position chrX:135,296,015 detected by DeepSAP and DRAGEN with novoSplice reports the event shifted 2 bases downstream.

Overall, the highest rates of true positive indels were found by novoSplice, DeepSAP, and DRAGEN with DeepSAP showing high sensitivity across datasets. Specific examples in the Baruzzo human T2 dataset revealed notable differences among aligners in their ability to detect indels within gene bodies and near splice sites. For instance, in the *SIK1* gene (chr21:44,836,563), DeepSAP, novoSplice, and DRAGEN all successfully identified a true single-nucleotide insertion of C following an A (Figure 6b). In contrast, a three-base deletion of CCA in *SPDYE3* (chr7:99,916,818) was uniquely detected by DeepSAP (Figure 6c). A more complex case was observed in the *HAL* gene (chr12:96,386,456), where a dinucleotide insertion of CC near the donor site was correctly reported by DeepSAP and novoSplice. In comparison, DRAGEN mischaracterized the event, reporting three insertions and a mismatch at the donor site, along with a false-positive deletion at the acceptor site (chr12:96,384,309). Other aligners failed to report this splice junction entirely, likely due to the presence of insertion events near both the donor and acceptor sites, which posed a challenge for accurate alignment (Figure 6d). In *MAP7D3* (chrX:135,296,015), both DeepSAP and DRAGEN correctly detected a GC insertion, while novoSplice reported the same insertion but shifted two bases downstream (Figure 6e).

## Discussion

We believe that DeepSAP marks a significant improvement in RNA-seq alignment, addressing long-standing challenges in splice junction detection, indel identification, and handling complex splicing patterns. The method demonstrates state-of-the-art performance, even on the most challenging benchmark datasets. By integrating TGGA GSNAP alignment with transformer-based deep learning models, DeepSAP excels in datasets characterized by high complexity. This combination of traditional alignment methods with cutting-edge machine learning substantially improves RNA-seq data analysis.

This improved performance stems from two key factors. First, it takes advantage of domain knowledge in the form of known transcripts through our implementation of TGGA GSNAP. In this method, a read is initially aligned, if possible, to the transcriptome, which is relatively straightforward because of the absence of intron gaps, and then remapped to the genome. If this alignment is not adequate, the read is then aligned to the genome using conventional splice-aware alignment methods that do allow for intron gaps. This dual approach yields higher accuracy and efficiency, especially at the ends of reads, where a genomic alignment may have insufficient evidence to determine the location of an intron, whereas known transcripts can often indicate its precise endpoints.

Second, the study utilizes fine-tuned transformer models for scoring splice junctions. These models capture intricate sequence patterns around splice donor and acceptor sites, significantly improving the accuracy of detecting splice junctions. We found that the DNABERT model provides superior performance compared with DNABERT2 and Nucleotide Transformer 2.5B models across various species and datasets, particularly in multi-species datasets such as MS150. Attention maps further reveal the critical role of distant exon and intron content in predicting splice sites accurately.

The methods of TGGA GSNAP and transformer-based models are highly complementary. GSNAP’s TGGA method demonstrates high sensitivity in identifying introns, particularly at the ends of reads, which can result in an increased rate of false positives. However, this is addressed by leveraging the transformer model’s capability to accurately predict false splice sites, thereby enhancing overall alignment accuracy. From the perspective of an aligner, enhanced sensitivity in intron detection inherently increases the likelihood of false positives. To mitigate this issue, incorporating a robust splicing score is essential. Traditionally, GSNAP has relied on an internal splice site scoring method using the local MaxEnt model(Yeo and Burge [2003]), which has proven effective. Nonetheless, modern transformer-based models provide superior accuracy in distinguishing true splice sites from false ones, offering a significant improvement over traditional approaches.

We have demonstrated the effectiveness of combining the two components of TGGA GSNAP and transformer models through our benchmarking studies with state-of-the-art RNA-seq aligners. In simulated datasets, DeepSAP consistently ranked as the top performer in recall and precision for read-level and base-level metrics. Its ability to detect complex splice junctions is particularly noteworthy in challenging datasets, demonstrating superior sensitivity compared with widely used aligners like STAR, HISAT2, DRAGEN, novoSplice and Subjunc. Its improved performance in both human and malaria datasets supports the generality of our strategy.

While aligners generally performed well on the human and malaria T1 and T2 datasets, the T3 datasets presented the greatest challenges, revealing significant differences in accuracy across aligners. The performance trends across various benchmarking metrics in the Baruzzo datasets indicate that as dataset complexity increases or species of origin shifts, aligner accuracy can vary significantly. Notably, DeepSAP stands out for its consistent accuracy across these conditions, demonstrating a strong capability in handling complex splice junction and transcriptomics variations in both human and malaria RNA-seq datasets. Although T3 dataset-level complexity is uncommon, it is frequently observed in cancer samples with elevated mutation burdens, particularly in metastatic cancers, mismatch repair-deficient tumors, and melanomas. These malignancies often demonstrate sequence variations comparable to the T3 dataset across their entire genomic landscape. Furthermore, while real-world samples may not exhibit overall T3-level complexity genome-wide, specific transcripts of interest within these samples may reach complexity levels comparable to those observed in the T3 dataset.

We believe that the accuracy of RNA-seq alignment will become increasingly important for personalized and individualized analyses of transcriptomes. In the past, analysis with existing aligners has been useful in exploring the shared, common repertoire of the transcriptome in the human population. But as sequencing becomes cheaper and more widely used, the next frontier will be to explore novel transcript phenomena in the presence of individual sequence variations and indels, where corroborating information from other individuals will be absent, and whose presence can be revealed only with the sensitivity and specificity provided by advances in RNA-seq alignment such as DeepSAP.

Nevertheless, after validating our strategy with the DeepSAP workflow, we are motivated to work on improving its speed, through approaches that include our own expertise in adapting and accelerating aligners by utilizing graphics processing units (GPUs). Another approach may include incorporating the transformer model to inform the alignment process rather than applying it as a post-processing step, which may also improve the overall accuracy.

## Methods

### Transcriptome-Guided Genomic Alignment

To leverage the domain knowledge contained in known transcripts, we have developed the methodology of TGGA GSNAP. Essentially, TGGA first aligns a given read to a transcriptome and, if such an alignment is unsuccessful, it subsequently attempts to align the read to the genome using conventional splice-aware alignment methods that accommodate intron gaps. Although this idea seems relatively straightforward, a successful implementation depends upon the development of specific methods for (1) performing accurate and efficient transcriptome alignment, (2) determining whether to accept transcriptome-based alignments or proceed to genome-based alignment, and (3) converting transcriptome alignments to genomic alignments, resolving instances where multiple transcript alignments are plausible but yield differing results when mapped onto the genome.

Before we can align RNA-seq reads, we need to pre-process the transcriptome and genome. We assume that a transcriptome is available as exon coordinates on a given genome, and such sequences are readily available in GFF or GTF formats from standard genome assemblies hosted by sites like NCBI and Ensembl. Alternatively, if transcript sequences are available without their genomic coordinates, these coordinates can be obtained by aligning the transcripts to a given genome using RNA long-read aligners such as GMAP.

For subsequent alignment, the transcriptome is pre-processed using the standard hash table format that GSNAP uses for genomes, with k-mers tabulated every 1 bp, in contrast to the default 3 bp used by GSNAP for large genomes. Additionally, the transcriptome sequence is stored in a linear format, similar to that used by GSNAP for genomes, with two bits for each nucleotide. The high bits and low bits are stored in separate arrays, and the bits are organized in 128-bit words to allow efficient SIMD (Single Instruction, Multiple Data) operations(Lemire et al. [2016]).

The genome is also pre-processed in the standard hash table format used by GSNAP, with k-mers at 3-bp intervals, and the nucleotide sequence in a linear format. A recent improvement in GSNAP involves using regional suffix arrays, where each array covers a non-overlapping 65,536-bp region of the genome. These arrays borrow upon the idea of local indexing introduced by HISAT, although their implementation is based upon the Burrows-Wheeler Transform and FM index, rather than suffix arrays. This region size was chosen to be represented by a suffix array with 16-bit elements, enabling the interrogation of one, two, or three regions to search for introns up to 200,000 bp, as typically observed in human transcripts. These regional suffix arrays help identify introns at the ends of reads during genomic alignment. However, since introns are not expected in transcriptome alignments, these regional suffix arrays are not necessary for TGGA. Finally, the exon coordinates for each transcript are stored for subsequent conversion of transcriptome alignments to genomic alignments.

Once the transcriptome and genomes have been pre-processed, RNA-seq reads can be aligned to the transcriptome and genome. Even though alignment to the transcriptome avoids the issue of intron gaps, it must still be able to handle mismatches and potentially small indels. Alignment to the transcriptome therefore involves three methods, each one tried in succession until a sufficiently good alignment is found. Sufficiency is determined by a threshold (default 5%) of mismatches or unaligned bases in the alignment between the read and the transcriptome. For paired-end reads, both ends must align to the same transcript in the correct orientation for the paired-end alignment to be considered sufficient. One advantage of transcriptome alignment is that it removes the possibility of alignment gaps due to introns, and so alignments are generally expected to extend to the ends of the read, making it possible to have an accurate evaluation based on unaligned bases.

The first transcriptome alignment method is called exact search, which is intended to be fast and handle the vast majority of reads. In this method, the first and last k-mer in the read are looked up in the hash table. Since the hash table is based on transcriptome k-mers in their forward orientations (or sense direction) only, the first and last k-mers of the reverse complement of the read are also looked up. Each hash table lookup yields a list of positions in the transcriptome for the corresponding k-mer. We can therefore find exact alignments to the genome, or even alignments with mismatches between the end k-mers by taking the intersection of the two lists of positions, adjusting for their expected distance in the transcript from the distance between the k-mers in the read. For each candidate position in the intersection, we then compare the read nucleotides against the transcript nucleotides to determine the number of matches and mismatches. Intersections are performed efficiently using a modified version of the SIMD algorithm by Lemire et al.(Lemire et al. [2016]).

The second method is an exhaustive one called prevalence search that looks for positions in the transcriptome with the most k-mers that match to the read. We count the number of matching k-mers by collecting lists of positions for each k-mer in the read, and then performing a multi-way merge of those positions, adjusting for their position in the read. The merged and adjusted positions allow us to count the number of matching k-mers for each transcriptome position. We can then identify those positions with the most matching k-mers. For reads with an indel, these positions should correspond to the longer segment, but a nearby segment within indel distance but with fewer matches should also appear in the list. Therefore, the exhaustive method should yield a series of segments that approximate the read alignment.

The third method is called extension search, which essentially follows the idea of seed-and-extend. In this method, seed k-mers are selected starting from both ends of the read, and tried in succession inward, with each seed evaluated by its longest matching extension in the transcriptome. Although extension could be defined as the longest contiguous stretch of matches, we have found that it is more effective to find a breakpoint in the alignment between primarily matches and mismatches. The trimming breakpoint is found by computing cumulative scores from the medial end, where a match receives +1 point, and a mismatch receives –3 points, identifying the position in the alignment with the maximum score. Seeds are extended iteratively in a breadth-first algorithm. When an extension ends for a given seed, we assume that a mismatch or indel was responsible, and we therefore resume the exploration of seeds for that branch in the search by skipping one position in the read. In a breadth-first algorithm, each branch in the search corresponds to the same number of continuous segments that match perfectly to the transcriptome. Search terminates when the segment reaches the other end of the read, and therefore the breadth-first search attempts to find solutions with the fewest number of matching segments. Because the algorithm is greedy, it is not guaranteed to find the optimal solution in terms of matching segments. Nevertheless, this algorithm works well when the exact method fails, because it handles cases where a mismatch occurs in either the first or last k-mer of the read, or when a read contains an indel. The extension method will therefore also yield a series of segments that approximate the read alignment.

Approximate alignments yielded by the exhaustive and extension methods are subsequently refined by procedures that identify the precise position of indels to minimize the number of mismatches. Resulting alignments from all methods are evaluated to see whether they satisfy the condition for ending the alignment process, or whether genomic alignment is called for. The condition for a paired-end read is finding a concordant alignment on the same transcript, whereas the condition for a single-end read is finding an alignment where 95% or more of the nucleotides on the read match those of a known transcript. Genomic alignment is especially necessary for reads corresponding to novel genes or isoforms that are not represented in the transcriptome.

Alignments from TGGA, whether they come from transcriptome-based or genome-based methods, are reported with respect to genomic coordinates. Therefore, a continuous alignment to a transcript may correspond to one or more exons in genome coordinates. The conversion of transcript coordinates to genome coordinates is performed using the known exon structure of the transcript on the genome. This procedure is generally straightforward, with the exception of indels that occur close (i.e., within 3 bp) to an exon-exon boundary in the transcript. In such cases, GSNAP uses a procedure that can resolve both the splice in the genome and the nearby indel to identify the precise location of both.

If the transcriptome-based methods do not find an alignment that is considered sufficient, then genome-based methods are used. These genome-based methods are similar to those for the transcript, corresponding to exact, extension, and exhaustive approaches to produce approximate alignments. For paired-end reads, these methods also post all genomic segments found during their computation, in order to facilitate paired-end alignment. If a concordant paired-end alignment cannot be found, a further search is made from the segments on each read to find any segments on the other read within a specified concordant distance, and an approximate alignment is constructed from those segments.

Approximate alignments are processed by procedures that both extend and refine the alignments. Extensions are found using both the genomic hash table and regional suffix arrays, which look for nucleotide sequences from the read that match the genome within a specified maximum indel or intron distance. Suffix arrays are searched using an exact method, which allows for no mismatches, or an exhaustive method using 4-mers, which does allow for mismatches. The extension methods can often yield multiple candidates for extending the alignment. These multiple candidates are handled differently depending on whether the unaligned end is on the inside or outside portion of a paired-end read. (For a single-end read, both ends are considered to be on the outside.) For the outside portion, if one candidate is clearly better in terms of a smaller intron length, that solution is used; otherwise the end is left unaligned. For the inside portion of a paired-end read, the candidates for one end are constrained by the genomic location of the other read. Among the allowable candidates, if one candidate is close enough to the other read, then that candidate is favored because we expect paired-end reads to come from a fragment with a relatively well-defined insert length. Procedures that refine approximate alignments use a splice model from MaxEnt to locate precise splice positions. Donor and acceptor splice junctions that appear at different positions in a read suggest the possibility of an indel, and such cases are handled by a procedure that solves for both a splice and nearby indel.

### Fine-Tuning Transformer Models

Traditional approaches to splice site prediction have relied on maximum entropy modeling(Yeo and Burge [2003]), machine learning(Pertea et al. [2001]; Xiong et al. [2015]), and, more recently, convolutional neural networks(Zuallaert et al. [2018]; Cheng et al. [2019]; Jaganathan et al. [2019]). These methods, however, face challenges in capturing the complex sequence patterns underlying splicing mechanisms(Riepe et al. [2021]; Smith and Kitzman [2023]). With the advancement of deep learning, genomics research is being revolutionized, introducing powerful new tools for analyzing genetic data. Among these advances, transformer-based LLMs have shown particular promise in decoding the complexities of splice site detection.

The transformer architecture is a groundbreaking neural network model for natural language processing(Vaswani et al. [2017]). Unlike traditional sequential models, transformers use self-attention mechanisms to process input sequences in parallel, capturing long-range dependencies between words. This architecture includes encoder and decoder components, employs multi-head attention, and position-wise feed-forward networks. The transformer achieved state-of-the-art performance on various NLP tasks, offering scalability and parallelization advantages. It marked a significant shift in the field, making it a foundational model in NLP and machine learning. In this study, we explore the application of cutting-edge transformer-based DNA foundation models, including DNABERT(Ji et al. [2021]), DNABERT2(Zhou et al. [2024]), and the Nucleotide Transformer(Dalla-Torre et al. [2023]), to the challenging task of splice site prediction.

DNABERT is a BERT-based(Devlin et al. [2018]) transformer model pre-trained on DNA sequences from the human reference genome. The sequences are tokenized into overlapping k-mers of lengths 3, 4, 5, or 6, with special tokens added at the beginning and end of each sequence, with maximum sequence length of 512 bases. DNABERT was pre-trained for 120,000 steps using a batch size of 2,000. During the first 100,000 steps, approximately 15% of the k-mers in each sequence were masked, as they could be trivially inferred from the neighboring k-mers. In the final 20,000 steps, the masking rate was increased to 20%(Ji et al. [2021]).

Building on the foundation of DNABERT, DNABERT2 introduces several key innovations to further improve genomic sequence modeling, using the same transformer encoder architecture but replacing k-mer tokenization with Byte Pair Encoding (BPE), a statistics-based data compression algorithm that constructs tokens by iteratively merging the most frequent co-occurring genome segments in the corpus. Additionally, DNABERT2 incorporates several recent advances in deep learning to enhance the model’s efficiency and performance: (i) Attention with Linear Biases (ALiBi)(Press et al. [2021]) replaces learned positional embeddings, allowing the model to overcome input length limitations. (ii) FlashAttention(Dao et al. [2022]), which speeds up the attention mechanism. (iii) Low-Rank Adaptation (LoRA)(Hu et al. [2021]), used during fine-tuning to enable parameter-efficient training. DNABERT2 was pre-trained on a multi-species genome dataset containing 32.49 billion nucleotide bases, covering genomes from 135 species(Zhou et al. [2024]).

Complementing these advances, the Nucleotide Transformer (NT) represents a collection of transformer-based DNA language models that have learned general nucleotide sequence representa-tions from 6kb unannotated genomic sequences, considering sequences of nucleotides as sentences and non-overlapping 6-mers as words. Pre-trained NT uses varying parameter sizes and datasets. (i) a 500 million parameter model pre-trained on sequences extracted from the human reference genome, (ii) a 500 million and (iii) a 2.5 billion parameter model both pre-trained on 3,202 genetically diverse human genomes, and (iv) a 2.5 billion parameter model based on 850 species including 11 model organisms. We chose the latest 2.5B parameters multi-species model to be tuned on classifying the splice junction sites since that model showed the highest normalized mean of MCC performance among the NT models for this specific downstream task(Dalla-Torre et al. [2023]).

For tuning DNABERT 6-mer, DNABERT2 and the 2.5B multi-species NT, we opted to generate our own splice-site datasets in a workflow that ensures transparency, customizability, reusability, and randomness. Each dataset consists of three files: train.csv for model tuning, dev.csv for evaluation during tuning, and test.csv for final validation, ensuring no data leakage between the files. Each of these CSV files contains (sequence, label) data points. Sequences are DNA sequences of specific lengths (90, 150, 200, 400) in which the splice donor or acceptor sites are in the center of the sequence. Labels indicate whether the sequence is donor (label=0), acceptor (label=1) or none (label=2). The negative data points were generated by (i) masking the genome around the annotated splice donor and acceptor sites, and (ii) mimicking splice junctions with varying intron sizes and splicing signals. The datasets were generated from (i) GRCh38 NCBI RefSeq assembly GCF_000001405.40 using transcripts with bio-types mRNA, lnc_RNA and transcript, (ii) GRCh38 GENCODE release 44 using all the annotated transcripts regardless of the bio-type and (iii) malaria ASM276v2. Furthermore, we tuned DNABERT using splice junctions with a window size of 150 from multi-species sources including the human RefSeq and GENCODE besides splice junctions from *Arabidopsis thaliana*, *Zea mays*, *Xenopus tropicalis*, *Saccharomyces cerevisiae*, *Drosophila melanogaster*, *Caenorhabditis elegans*, *Rattus norvegicus*, *Mus musculus*, *Plasmodium falciparum*, *Plasmodium vivax*, *Plasmodium berghei*, *Plasmodium knowlesi* The generated fine-tuning datasets are named as RefSeq90, RefSeq150, RefSeq200, RefSeq400, GENCODE150, GENCODE400 and MultiSpecies150.

### Applying Transformer Score to BAM Records

DeepSAP is designed for aligning short-read RNA-seq samples using the TGGA GSNAP aligner, followed by a transformer-based splice junction scoring scheme as a post-alignment step. It accepts short-read RNA-seq data in FASTQ format along with a fine-tuned transformer model as input, generating a BAM file after alignment using TGGA GSNAP and applying the scoring scheme based on the transformer’s prediction probabilities. The scoring is calculated as follows:

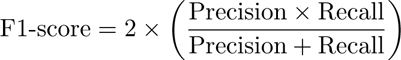

After generating the BAM file with GSNAP, DeepSAP performs two passes over the file. In the first pass, it identifies splice junctions in spliced reads, creating labeled sequences for processing by the transformer model. In the second pass, DeepSAP applies the transformer-based splice junction score to achieve several objectives: (i) soft-clipping novel splice junctions with low scores; (ii) adjusting MAPQ (Mapping Quality) values for multi-mapped reads containing novel splice junctions; and (iii) soft-clipping junctions where flanking bases have low Phred quality scores.

### Establishing Benchmarking Study With Simulated Datasets

We conducted a large-scale RNA-seq alignment benchmarking study using simulated datasets from two origins. First, datasets were obtained from(Baruzzo et al. [2017]) which include simulated RNA-seq datasets from human and malaria with varying levels of complexities. The least complex dataset, T1, contained low polymorphism rate (0.001 substitution, 0.0001 indels with 0.005 sequencing error rate) followed by T2, which contained moderate polymorphism rate (0.005 substitution, 0.002 indels and 0.01 sequencing error rate). Finally, T3 is the most complex dataset in the study with polymorphism rate (0.03 substitution, 0.005 indels and 0.02 sequencing error rate). Secondly, we also simulated datasets using the SimBA tools suite(Audoux et al. [2017]) with similar levels of complexities of the previous datasets and using the same genomes and annotations. The twelve datasets were then aligned using state-of-the-art RNA-seq aligners including DeepSAP with TGGA GSNAP v24.06.2024 and DNABERT MS150, Illumina’s DRAGEN™ v4.0.3, novoSplice v0.8.4(Berakdar et al. [2019]), STAR v2.7.10a(Dobin et al. [2013]), HISAT2 v2.2.1(Kim et al. [2019]) and Subjunc v2.0.1(Liao et al. [2013]). To assist the performance of the RNA-seq aligners, we compared the alignment output for each aligner with the known ground truth provided by the simulators. For Baruzzo datasets, we used the original methods in calculating different levels of benchmarking metrics as introduced in the Baruzzo study and we added F1 score calculated as follows:

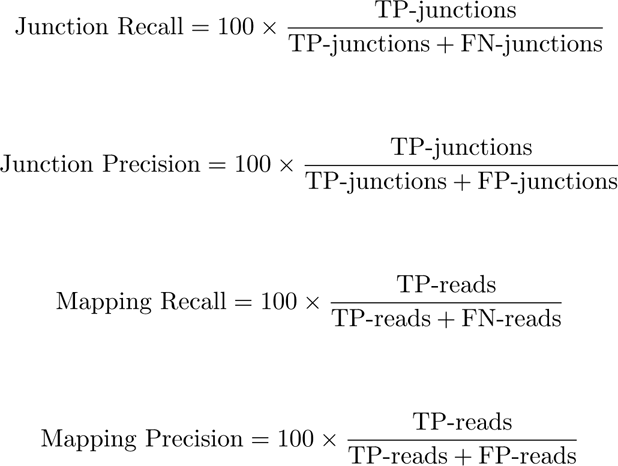

The benchmarking results are organized as a Nextflow pipeline that runs inside a docker container to allow for maximum reproducibility and transparency. After the Nextflow run is completed, the benchmarking results are visualized in a ShinyApp that is publicly accessible, Figure S1.

### Detection of Complex Splice Junctions

To closely evaluate how well the RNA-seq aligners in our benchmark recovered splice junctions com-pared to the ground truth, we analyzed the Baruzzo human and malaria simulated datasets. Splice junction counts were calculated by processing the sorted aligned BAM files from each aligner using fea-tureCounts(Liao et al. [2014]) with the following options: (-J -O -Q 0 -t exon -g gene_id -p -M), which enabled the inclusion of overlapping features and multi-mapped reads, as well as proper handling of paired-end data.

Detected junctions were then compared with the ground truth using scatter plots generated in Python with the *matplotlib* library. For aligner-to-truth comparisons, we applied the Theil–Sen robust linear estimator using the *theilslopes* function from SciPy with default parameters. For pairwise aligner comparisons, we used the Pearson correlation coefficient via SciPy’s *pearsonr* function to assess similarity in junction detection profiles, reporting both correlation values and associated p-values. These analyses revealed each aligner’s ability to recover complex splicing patterns, with particular emphasis on the challenging T3 dataset, which includes simulated indels and complex splice junction.

To complement our analysis on simulated datasets, we extended the evaluation to real RNA-seq data using samples from a glioma study by Siddaway et al.(Siddaway et al. [2022]). However, in the absence of a definitive ground truth for splice junctions in real datasets, DeepSAP’s splice junction detection performance was assessed by comparing its results with those of other aligners using concordance scatter plots. To estimate the abundance of splice junctions Feature-Counts(Liao et al. [2014]) was employed on each aligner’s sorted BAM files, with the following options: (-J -O -Q 0 -t exon -g gene_id -p -M) allowing both overlapping features and multi-mapped reads. The concordance of known and novel splice junctions across aligners was then assessed by visualizing them in scatter plots. These scatter plots (Figure 5a) highlight DeepSAP’s concordance with other aligners in splice junction detection. Points near the diagonal represent junctions with similar counts to those reported by DeepSAP for the same site. Points along the line x = 1 correspond to junctions uniquely identified by DeepSAP, while points along y = 1 represent junctions detected exclusively by other aligners. Counts are shown on a log scale using log(count + 1) to accommodate a wide range of values while avoiding issues with zero counts.

To ensure reproducibility, the entire analysis was managed within a Snakemake(Köster and Rahmann [2012]) pipeline, streamlining the workflow across aligners and handling all alignment and analysis steps within a reproducible framework.

### Detecting Insertions and Deletions

To assess the ability of DeepSAP to detect indels in RNA-seq data, we compared its performance against several widely used RNA-seq aligners using the simulated datasets from Baruzzo et al.(Baruzzo et al. [2017]). We used all aligners from the benchmarking study with their respective default settings to ensure compatibility with indel detection and accurate splice junction mapping.

To identify indels from each aligner’s output BAM file, we implemented a standardized variant calling and indel normalization workflow. Aligned BAM files were first converted into pileup format using samtools *mpileup*, subsequently small insertions and deletions were detected using VarScan’s *mpileup2indel* function with default parameters. To enable consistent and fair comparison of variant calls across aligners, we applied bcftools *norm* with parameters (-m -any) to left-align and normalize multi-allelic indels, followed by bcftools *sort* to ensure proper VCF record ordering. Finally, we used the bcftools *isec* tool to compute intersections between aligner-derived VCFs and the ground truth provided by Baruzzo et al. This pipeline ensured alignment-independent variant comparison by enforcing consistent indel representation across aligners. Performance was quantified using true positives (TP), false positives (FP), and false negatives (FN), with percentages reported for each metric to reflect each tool’s ability to accurately recover indels and minimize erroneous calls.

## Data Availability

The human reference genome used in this study is GRCh37 and is available at https://ftp.ebi.ac.uk/pub/databases/gencode/Gencode_human/release_40/GRCh37_mapping/ GRCh37.primary_assembly.genome.fa.gz. The human annotation, as well as the malaria reference genome and annotation, along with human and malaria T1, T2, and T3 datasets are available from the Baruzzo homepage http://bioinf.itmat.upenn.edu/BEERS/bp1/datasets.html. However, the SimBA datasets are simulated in-house using SimCT. For further details on the simulation of SimBA datasets, and the genome reference and annotation files used for fine-tuning the transformer model, please refer to the supplemental document Nextflow.md found in the accompanying **GitHub repository:** https://github.com/clara-parabricks-workflows/DeepSAP.

## Code Availability

GSNAP is available open-source under the Apache 2.0 license; past and recent versions, including the 2024-06-24 version used in this study, can be downloaded from http://research-pub.gene.com/gmap. The DeepSAP pipeline can be run within a Docker container, which is available at **NGC Catalog** https://catalog.ngc.nvidia.com/orgs/nvidia/teams/clara/containers/clara-parabricks-deepsap using version tag v0.0.1.

The supplemental files include detailed documentation and scripts related to this study, hosted in a public **GitHub repository:** https://github.com/clara-parabricks-workflows/DeepSAP.A com-prehensive overview of these files can be found in the manuscript_data_code/README.md of the repository. Additionally, to explore the RNA-seq aligner benchmarking results, we have visu-alized the data in an interactive Shiny app dashboard, accessible at **DeepSAP-Dashboard:** https://rna-seqbenchmark.shinyapps.io/DeepSAP-Dashboard

## Authors Information

### Contributions

P.V, T.D.W and F.B conceived the research idea. PV, T.D.W, F.B, T.Z., and M.S. provided critical feedback and helped in drafting the manuscript. P.V and F.B. developed the benchmarking pipelines, provided feedback on improving TGGA GSNAP and performed fine-tuning and performance evaluation of transformer models. T.D.W. developed TGGA GSNAP and PV, F.B. developed the post-alignment step. PV, T.D.W and F.B performed data analysis, interpretation and figure generation. P.V and T.D.W provided advice on study design. P.V supervised this work. All authors reviewed and approved the manuscript draft.

### Disclosure

The authors declare the following competing interests:

Fadel Berakdar, Tong Zhu, Mehrzad Samadi, and Pankaj Vats are salaried employees of **NVIDIA Corporation**, a leading technology company renowned for its pioneering work in graphics processing units (GPUs), artificial intelligence (AI), and high-performance computing.

Thomas D. Wu is a salaried employee of **Genentech**, a member of the Roche group, which develops and markets drugs for profit.

## Notes

https://ftp.ebi.ac.uk/pub/databases/gencode/Gencode_human/release_40/GRCh37_mapping/GRCh37.primary_assembly.genome.fa.gz

http://bioinf.itmat.upenn.edu/BEERS/bp1/datasets.html

https://github.com/clara-parabricks-workflows/DeepSAP

http://research-pub.gene.com/gmap

https://catalog.ngc.nvidia.com/orgs/nvidia/teams/clara/containers/clara-parabricks-deepsap

https://rna-seqbenchmark.shinyapps.io/DeepSAP-Dashboard

### Competing Interest Statement

The authors declare the following competing interests:
Fadel Berakdar, Tong Zhu, Mehrzad Samadi, and Pankaj Vats are employees of NVIDIA, a leading technology company renowned for its pioneering work in graphics processing units (GPUs), artificial intelligence (AI), and high-performance computing.
Thomas D. Wu is an employee of Genentech, a member of the Roche group, which develops and markets drugs for profit.

