## Supplemental_figures for "DeepSAP: Improved RNA-Seq Alignment by Integrating Transcriptome Guidance with Transformer-Based Splice Junction Scoring"

[1]NVIDIA Corporation, 2788 San Tomas Expy, Santa Clara, 95051, CA, USA.
[2]Genentech, Inc, 1 DNA Way, South San Francisco, 94110, CA, USA.

†These authors contributed equally to this work.

### Supplemental Figures and Tables

#### Contents

**Supplemental Figure 1: Nextflow Pipeline for Benchmarking RNA-seq Aligners** **2**

**Supplemental Figure 2: Comparing DNABERT MS150 and RefSeq150 Models** **3**

**Supplemental Figure 3: Benchmarking RNA-seq Aligners Using SimBA** **4**

**Supplemental Figure 4: Detection of Complex Splice Junctions in Baruzzo Datasets** **5**

**Supplemental Figure 5: Detection of Complex Splice Junctions in SRR6781181 Dataset** **6**

**Supplemental Table 1: Benchmarking Metrics for Baruzzo Datasets** **7**

### Supplemental Figure 1: Nextflow Pipeline for Benchmarking RNA-seq Aligners

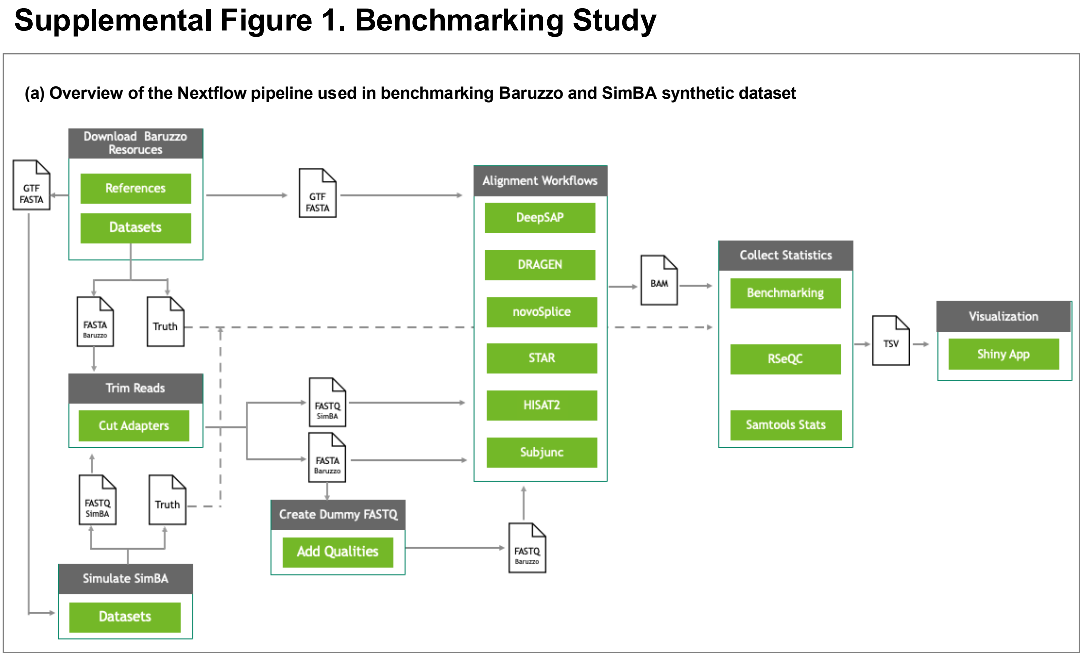


**Supplemental Figure 1:** Overview of the benchmarking study utilizing the Baruzzo and SimBA datasets, managed within a Nextflow pipeline. The pipeline automates the process, including downloading reference genomes, datasets, and reads, followed by read trimming. Additionally, Baruzzo FASTA read files are converted into FASTQ format with a fixed quality score of Q=25 since some aligners only accept FASTQ input. Once preprocessing is complete, the Nextflow pipeline executes alignment jobs using default settings to generate BAM files. These files are then analyzed by the Baruzzo and SimBA benchmarking scripts, ultimately producing a results.tsv file, which can be visualized in an interactive Shiny app.

### Supplemental Figure 2: Comparing DNABERT MS150 and RefSeq150 Models

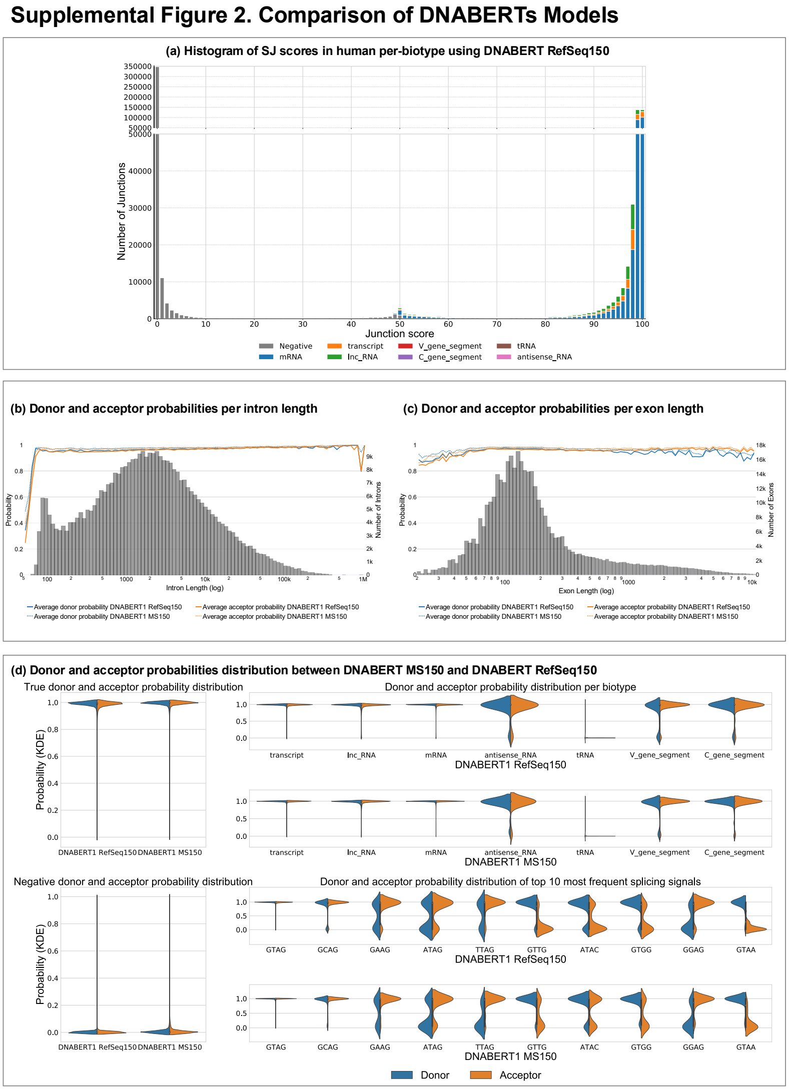


**Supplemental Figure 2:** (a) Histogram of SJ scores of different transcript biotypes in RefSeq human dataset predicted by DNABERT RefSeq150. (b) Donor and acceptor scores across intron lengths for DNABERT MS150 and DNABERT RefSeq150. (c) Donor and acceptor scores across exon lengths for DNABERT MS150 and DNABERT RefSeq150. (d) Distribution of donor and acceptor probabilities across various biotypes and splicing signals in human RefSeq splice junctions for DNABERT MS150 and DNABERT RefSeq150.

### Supplemental Figure 3: Benchmarking RNA-seq Aligners Using SimBA

Supplemental Figure 3. SimBA Benchmarking Results

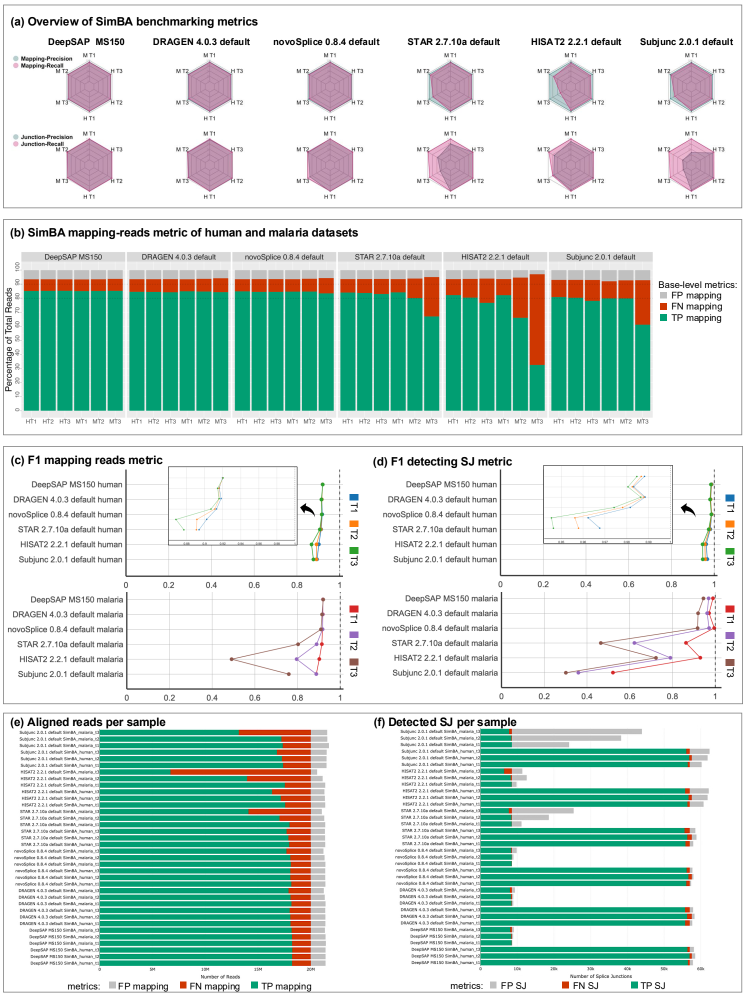


**Supplemental Figure 3:** (a) Overview of benchmarking results using the SimBA dataset. DeepSAP, DRAGEN, and novoSplice demonstrate consistent performance across various metrics, species, and complexity levels. (b) Percentage distribution of SimBA read mapping metrics across different aligners and datasets. (c) F1-score for the SimBA read mapping metric in human and malaria datasets, with a zoomed-in plot highlighting differences in F1-score ranges close to 1. (d) F1-score for the SimBA read junction metric in human and malaria datasets, with a zoomed-in plot highlighting differences in F1-score ranges close to 1. (e) Bar plots showing the number of aligned reads per sample. DeepSAP achieves high true-positive rates (green bars) across samples. (f) Bar plots illustrating the number of detected splice junctions per sample. DeepSAP exhibits a high true-positive rate (green bars) across samples.

### Supplemental Figure 4: Detection of Complex Splice Junctions in Baruzzo Datasets

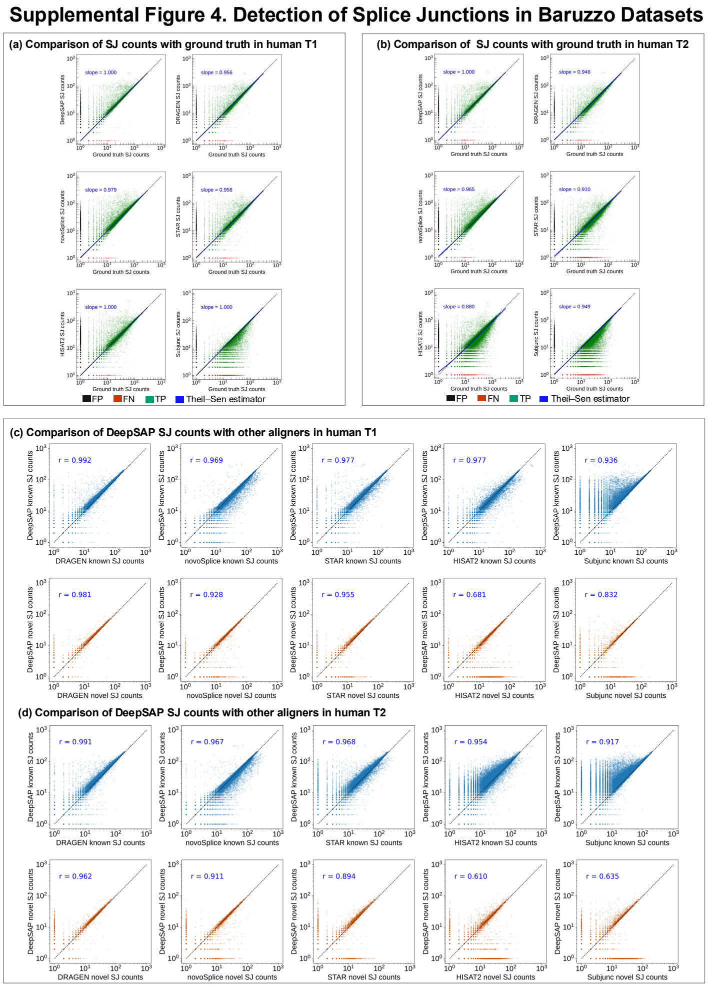


**Supplemental Figure 4:** (a) Splice junction counts from each aligner are compared against the ground truth in the Baruzzo human T1 dataset. The axes are displayed on a logarithmic scale using log(count + 1) to avoid log(0) values. The Theil-Sen slope (blue line) was computed to estimate the linear trend between each aligner's splice junction counts and the ground truth. DeepSAP demonstrates a near-perfect Theil-Sen slope, alongside HISAT2 and Subjunc, while other aligners also perform well in this dataset. (b) Splice junction counts from each aligner are compared against the ground truth in the Baruzzo human T2 dataset. The Theil-Sen slope (blue line) was computed to estimate the linear trend. DeepSAP achieves the best Theil-Sen slope among the aligners in this dataset. (c) Pairwise comparisons of splice junction counts between DeepSAP and other aligners in the Baruzzo human T1 dataset are shown. The axes are plotted on a logarithmic scale using log(count + 1) to avoid log(0) values. Known splice junctions are represented in blue, while novel junctions are in orange. DeepSAP shows strong correlation with statistically significant p-values when compared with DRAGEN, novoSplice, and STAR for both known and novel junctions. (d) Pairwise comparisons of splice junction counts between DeepSAP and other aligners in the Baruzzo human T2 dataset are presented. The axes are displayed on a logarithmic scale using log(count + 1) to avoid log(0) values. Known splice junctions are shown in blue, and novel junctions in orange. DeepSAP demonstrates a strong correlation with statistically significant p-values when compared with DRAGEN and novoSplice for both known and novel junctions.

### Supplemental Figure 5: Detection of Complex Splice Junctions in SRR6781181 Dataset

**Supplemental Figure 5. Detection of Splice Junctions in Real Datasets**

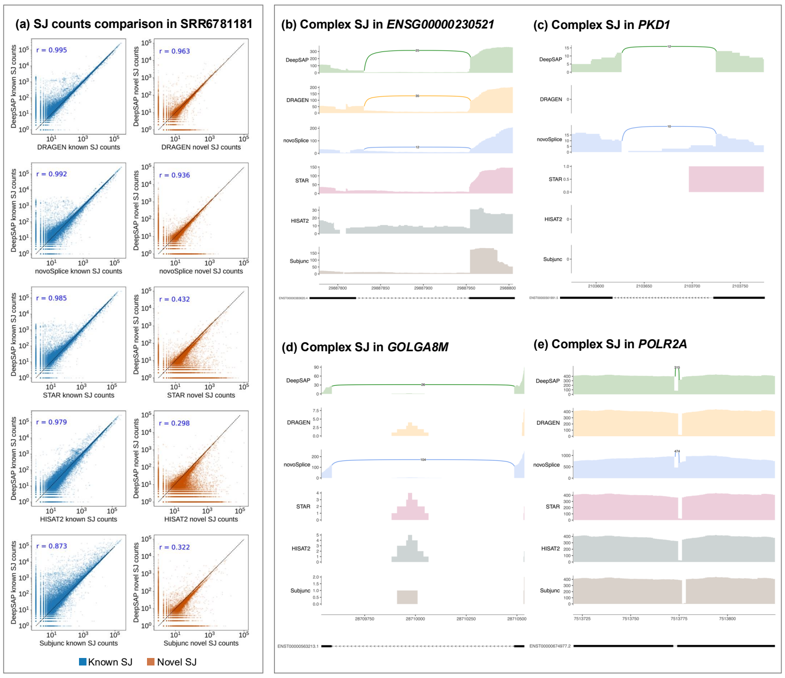


**Supplemental Figure 5:** (a) Comparison of splice junctions between DeepSAP and other aligners in the SRR6781181 dataset. The axes are displayed on a logarithmic scale using log(count + 1) to avoid log(0) values. Known splice junctions are shown in blue, while novel junctions are represented in orange. DeepSAP exhibits a high Pearson correlation with DRAGEN and novoSplice, with statistically significant p-values for both known and novel junctions. (b) A splice junction in ENSG00000230521, identified by DeepSAP, DRAGEN, and novoSplice but undetected by other aligners. (c) A splice junction in *PKD1* gene, detected by DeepSAP and novoSplice but missed by other aligners. (d) A splice junction in *GOLGA8M*, identified by DeepSAP and novoSplice but not by other aligners. (e) A splice junction in *POLR2A*, detected by DeepSAP and novoSplice but absent in other aligners.

### Supplemental Table 1: Benchmarking Metrics for Baruzzo Datasets

| Run | Dataset | F1 Junction | F1 Skipping | F1 Insertion | F1 Deletion |
|---|---|---|---|---|---|
| DeepSAP MS150 | Human T1 | **98.846** | **98.965** | **90.126** | **91.935** |
| DRAGEN 4.0.3 default | Human T1 | 95.794 | 96.118 | 82.791 | 84.779 |
| novoSplice 0.8.4 default | Human T1 | 94.738 | 95.361 | 86.032 | 87.095 |
| STAR 2.7.10a default | Human T1 | 95.949 | 95.823 | 75.273 | 82.561 |
| HISAT2 2.2.1 default | Human T1 | 97.099 | 97.455 | 78.073 | 79.999 |
| Subjunc 2.0.1 default | Human T1 | 97.092 | 94.823 | 85.884 | 88.705 |
| DeepSAP MS150 | Human T2 | **98.591** | **98.609** | **87.773** | **89.338** |
| DRAGEN 4.0.3 default | Human T2 | 95.488 | 95.698 | 80.919 | 82.904 |
| novoSplice 0.8.4 default | Human T2 | 94.234 | 94.896 | 83.617 | 84.061 |
| STAR 2.7.10a default | Human T2 | 93.352 | 93.042 | 66.848 | 74.457 |
| HISAT2 2.2.1 default | Human T2 | 90.844 | 90.996 | 56.991 | 58.196 |
| Subjunc 2.0.1 default | Human T2 | 93.053 | 90.423 | 81.281 | 83.437 |
| DeepSAP MS150 | Human T3 | **93.970** | **93.822** | **67.546** | **68.848** |
| DRAGEN 4.0.3 default | Human T3 | 88.619 | 87.552 | 59.385 | 60.293 |
| novoSplice 0.8.4 default | Human T3 | 85.413 | 86.178 | 58.508 | 59.775 |
| STAR 2.7.10a default | Human T3 | 57.093 | 54.236 | 23.566 | 26.753 |
| HISAT2 2.2.1 default | Human T3 | 10.720 | 10.517 | 1.744 | 1.685 |
| Subjunc 2.0.1 default | Human T3 | 41.107 | 39.448 | 22.123 | 21.389 |
| DeepSAP MS150 | Malaria T1 | **99.529** | **99.226** | 89.538 | 89.400 |
| DRAGEN 4.0.3 default | Malaria T1 | 95.487 | 95.069 | 81.039 | 80.903 |
| novoSplice 0.8.4 default | Malaria T1 | 96.545 | 96.384 | 89.398 | 88.913 |
| STAR 2.7.10a default | Malaria T1 | 96.710 | 39.208 | 78.226 | 83.120 |
| HISAT2 2.2.1 default | Malaria T1 | 96.090 | 61.935 | 75.593 | 75.996 |
| Subjunc 2.0.1 default | Malaria T1 | 95.048 | 2.038 | **90.971** | **91.370** |
| DeepSAP MS150 | Malaria T2 | **99.327** | **98.871** | **86.617** | **87.199** |
| DRAGEN 4.0.3 default | Malaria T2 | 95.227 | 91.598 | 78.389 | 79.694 |
| novoSplice 0.8.4 default | Malaria T2 | 96.299 | 96.148 | 85.881 | 86.050 |
| STAR 2.7.10a default | Malaria T2 | 94.365 | 9.169 | 71.065 | 77.227 |
| HISAT2 2.2.1 default | Malaria T2 | 89.469 | 43.924 | 55.754 | 55.858 |
| Subjunc 2.0.1 default | Malaria T2 | 88.899 | 1.212 | 85.724 | 87.010 |
| DeepSAP MS150 | Malaria T3 | **95.279** | **92.728** | **67.454** | **66.591** |
| DRAGEN 4.0.3 default | Malaria T3 | 88.481 | 51.759 | 54.845 | 53.658 |
| novoSplice 0.8.4 default | Malaria T3 | 87.530 | 87.268 | 60.330 | 58.633 |
| STAR 2.7.10a default | Malaria T3 | 62.839 | 0.379 | 32.939 | 34.411 |
| HISAT2 2.2.1 default | Malaria T3 | 10.820 | 7.781 | 1.626 | 1.567 |
| Subjunc 2.0.1 default | Malaria T3 | 33.311 | 0.140 | 24.853 | 22.845 |

**Supplemental Table 1:** F1 scores for various Baruzzo base-level metrics across human and malaria datasets. DeepSAP consistently achieves top performance, with the only exception being insertions and deletions in the malaria T1 dataset, where it performs closely to the top aligner, Subjunc.
